# Excitation–inhibition balance controls coupling stability and network reorganization in a plastic Kuramoto model

**DOI:** 10.1101/2024.06.17.599234

**Authors:** Satoshi Kuroki, Kenji Mizuseki

## Abstract

Sleep and rest, characterized by synchronized neuronal activity that emerges under large shifts in the excitation–inhibition balance, are crucial for synaptic reorganization in the brain. However, the network dynamics that permit rewiring without erasing stable connections remain unclear. To address this question, we extended a Kuramoto framework with excitation–inhibition balance by adding Hebbian and homeostatic plasticity. The resulting model showed that networks with robust inhibition consistently exhibited desynchronized dynamics and stable couplings. In contrast, networks with weaker inhibition exhibited a bistable regime, in which couplings of intermediate strengths fluctuated while strong couplings remained stable. These findings suggest that the dynamic interplay between network activity and plasticity selectively stabilizes stronger connections while permitting flexible reorganization of weaker ones, providing a potential mechanism for network reorganization during sleep and rest.

## Introduction

Empirical evidence suggests that brain activity during sleep and rest promotes memory consolidation, knowledge abstraction, and creative insight ^1,2^ through the reorganization of neural circuits ^3,4^. Such reorganization should rely on mechanisms distinct from those involved in learning during task performance. Indeed, synaptic plasticity during sleep and rest improves task performance ^5,6^.

Neuronal synchronization occurs in two distinct states. Collective neuronal firing is desynchronized during active wakefulness, including task engagement, movement, and attentional states, as well as during rapid eye movement (REM) sleep, but more synchronized during quiet wakefulness and non-REM (NREM) sleep. Throughout the paper, we refer to these regimes descriptively as wake-like/desynchronized and sleep/rest-like/synchronized or bistable states, depending on the model dynamics being discussed. The transition between these states is associated with shifts in the excitation–inhibition (EI) balance. Firing rates of inhibitory neurons and synaptic conductance of inhibitory synapses are significantly elevated during wake-like states, while those of excitatory synapses show comparatively smaller changes ^7–10^. These observations raise the possibility that, during sleep/rest-like states, EI-balance-dependent changes in collective neuronal synchronization interact with synaptic plasticity to reshape network structure. However, detailed mechanisms by which neuronal synchronization in sleep/rest-like states reshapes network structure through plasticity remain unclear.

Collective synchronization of oscillators, including neurons, has been studied using coupled-oscillator models. Among these, the Kuramoto model is one of the simplest frameworks^11,12^, and numerous variants have been proposed. One variant, the EI-Kuramoto model, incorporates the EI balance by dividing oscillators into excitatory and inhibitory groups and assigning four interaction types between them^13,14^. We have previously shown that tuning these four strengths toggles the network between synchronized, desynchronized, and bistable states, potentially reflecting a neuronal synchronization mechanism^14^. That study, however, kept all coupling weights—defined in this paper as pair-specific interaction strengths—fixed and identical, and therefore could not address how EI-balance-dependent dynamical states shape synaptic coupling through plasticity over time.

In this study, we focused on Hebbian and homeostatic plasticity. Hebbian plasticity, based on the principle “fire together, wire together,” strengthens synapses between neurons that fire in close temporal proximity^15^. In contrast, homeostatic plasticity regulates neuronal excitability and synaptic strength to maintain network stability, thereby preventing excessive excitation or quiescence^16^. The interplay between these plasticity mechanisms enables the emergence of diverse neuronal functions and firing patterns^17–20^. Previous studies have incorporated plasticity into Kuramoto models, enabling coupling weights to adapt based on the oscillator activity to examine their impact on system synchronization^21– 31^. However, these studies focused on promoting synchronization under a single dynamic state. How distinct collective dynamics, controlled by the EI balance, differentially shape synaptic plasticity remains unexplored.

To address these gaps, we developed a plastic EI (pEI)-Kuramoto model in which excitatory–excitatory couplings undergo Hebbian potentiation and homeostatic regulation. We found that strong inhibition produced a desynchronized regime with relatively stable couplings, whereas weaker inhibition produced a bistable regime in which intermediate-strength couplings fluctuated while strong couplings remained stable. These results suggest that EI-balance-dependent collective dynamics selectively stabilize strong couplings while permitting flexible reorganization of weaker ones, providing a potential mechanism for network reorganization during sleep and rest.

## Results

### Establishment of the pEI-Kuramoto model

The pEI-Kuramoto model comprised two oscillator groups: excitatory units, referred to as excUnits, and inhibitory units, referred to as inhUnits. These specific terms clarify that the units represented excitatory and inhibitory factors in this model, rather than actual neurons or synapses. Four interaction strengths were assigned between the groups, with only excitatory-excitatory couplings (*J*_*ee*_) being distributed and plastic. The other interaction strengths, *K*_*ii*_ (repulsive strength between inhUnits), *K*_*ei*_ (attractive strength from excUnits to inhUnits), and *K*_*ie*_ (repulsive strength from inhUnits to excUnits) were constant across time and across unit pairs (Figure 1A; see Materials and Methods). The interactions are modeled as zero-phase-lag sine functions. The interactions from excUnits are purely attractive, while those from inhUnits are purely repulsive. Accordingly, the terms “excitatory” and “inhibitory” are used synonymously with “attractive” and “repulsive,” respectively.

**Figure 1.**
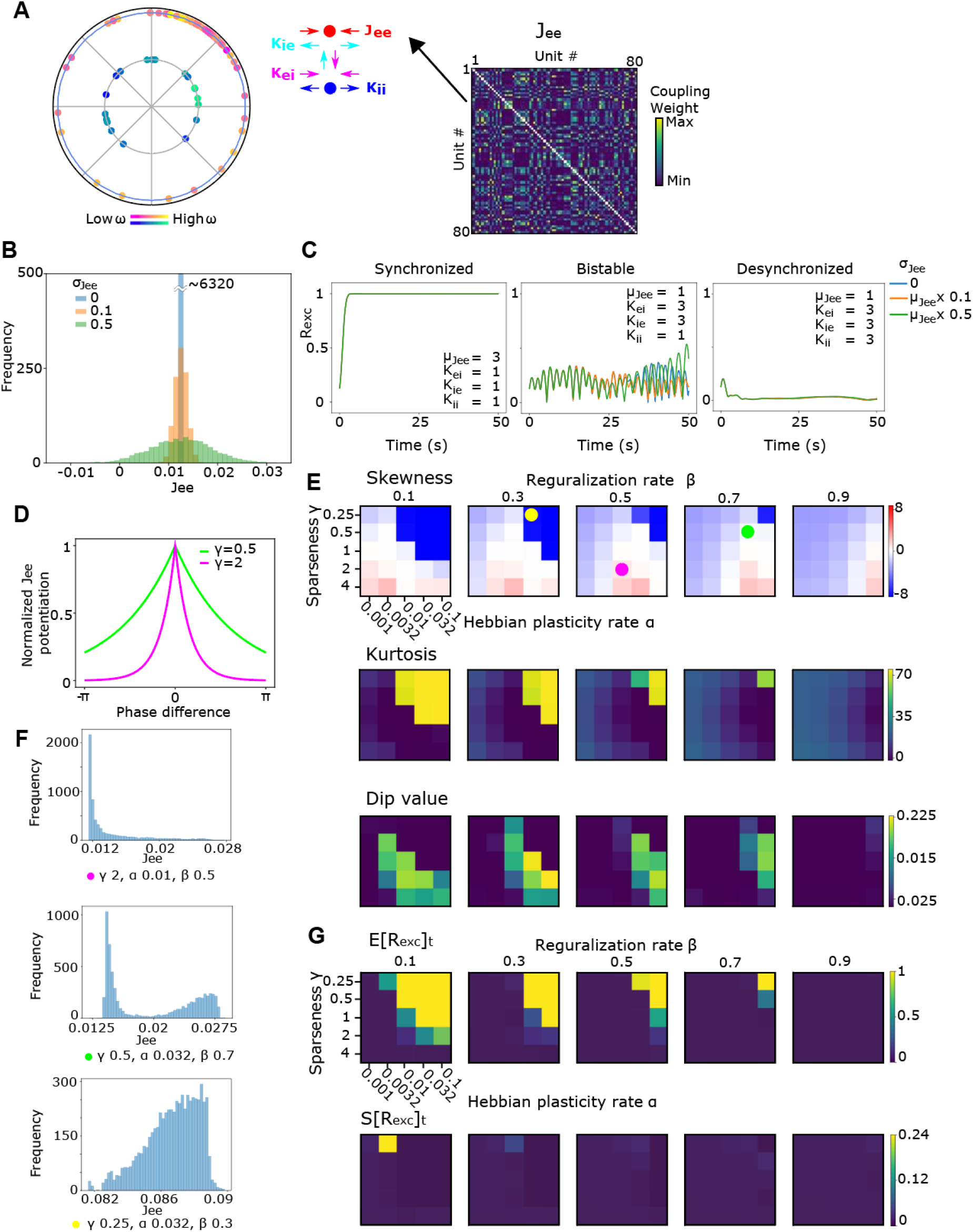
EI-Kuramoto model with distributed *J*_*ee*_ and plasticity. (A) Schematic representation of the pEI-Kuramoto model. Left: Oscillator units are excitatory (warm-colored) or inhibitory (cool-colored), each with its own natural frequency (shown by color gradients). Middle: *J*_*ee*_ functions attractively between excUnits (red arrows). *K*_*ei*_ is the attractive interaction strength from excUnit to inhUnit (magenta arrow). *K*_*ii*_ represents the repulsive strength between inhUnits (blue arrows). *K*_*ie*_ is the repulsive strength from inhUnit to excUnit (cyan arrow). *K*_*ei*_, *K*_*ii*_, and *K*_*ie*_ are constants that are applied uniformly across all units. Right: Each *J*_*ee*_ unit pair has its own coupling weight and undergoes plastic modification. (B) Distribution of *J*_*ee*_ coupling weights with different *σ*_*Jee*_. *μ*_*Jee*_ = 1, *N*_*exc*_ = 80. (C) *R*_*exc*_ time traces of synchronized, bistable, and desynchronized states with each *σ*_*Jee*_ and no plasticity. (D) Exponential rules of Hebbian potentiation plasticity. The *J*_*ee*_ weight of each pair of units is potentiated based on the phase difference between the units. (E) Evaluation of the *J*_*ee*_ distribution at the last frame (*t* = 50 s) with various plasticity parameters. Skewness (top panels), kurtosis (middle), and dip values (bottom). (F) *J*_*ee*_ distribution of exponential Hebbian plasticity rules at the last frame (*t* = 50 s) with various parameter settings. The parameter settings of the top, middle, and bottom panels correspond to those marked by magenta, green, and yellow circles in (E). (G) Evaluation of state synchrony with various plasticity parameters. E[*R*_*exc*_]_t_ and S[*R*_*exc*_]_t_ denote the time-averaged mean and standard deviation, respectively, of the order parameter for excUnits, *R*_*exc*_. The parameters used for evaluation were *N*_*exc*_ = 80, *N*_*inh*_ = 20, *μ*_*Jee*_ = 1, *σ*_*Jee*_ = 0.1, *K*_*ei*_ = 3, *K*_*ie*_ = 3, *K*_*ii*_ = 3, corresponding to a desynchronized state of the system without plasticity (E–G).

First, we investigated how the distribution of pair-specific excitatory-excitatory coupling weights *J*_*ee*_ affected the dynamics of the pEI-Kuramoto model in the absence of plasticity (Figure 1B and C). In our previous study, we demonstrated that the Kuramoto model with EI balance (EI-Kuramoto model) exhibits three distinct dynamic states—synchronized, bistable, and desynchronized—depending on the balance between the constant values of the interaction strengths^14^. In this study, *J*_*ee*_ followed a Gaussian distribution with a mean of *μ*_*Jee*_ and a standard deviation of *σ*_*Jee*_, where *σ*_*Jee*_ was set to 0, 1/10, and 1/2 of *μ*_*Jee*_ (Figure 1B). The collective synchronization was quantified using the order parameter *R*, which represents the magnitude of the average vector between the oscillator units. In this simulation, regardless of the standard deviation value, all systems with a distributed *J*_*ee*_ also exhibited synchronized (high *R*), bistable (alternating high and low *R*), and desynchronized (low *R*) dynamics, as observed with constant (*σ*_*Jee*_ = 0) values (Figure 1C).

Next, we incorporated plasticity into *J*_*ee*_ by applying Hebbian potentiation and homeostatic regularization plasticity rules (see Materials and Methods). Among the various methods available for implementing Hebbian plasticity in the Kuramoto model, we initially selected the simplest method: the cosine function (Figure S1A; see Materials and Methods)^21–23,26,30,31^. In this process, when a pair of excUnits exhibits closer phases, their *J*_*ee*_ coupling weight is potentiated. Simultaneously, the *J*_*ee*_ distribution was rescaled to maintain the initial mean and standard deviation, thereby preventing overpotentiation through homeostatic regularization plasticity (see Materials and Methods). The system dynamics were evaluated using the parameter set (*μ*_*Jee*_ = 1, *σ*_*Jee*_ = 0.1, *K*_*ei*_ = 3, *K*_*ie*_ = 3, *K*_*ii*_ = 3), at which the system exhibited a desynchronized state in the absence of *J*_*ee*_ plasticity. Under the cosine rule, unit pairs with smaller phase differences acquire a larger *J*_*ee*_ weight. This rule causes the J_ee_ distribution, which was initially Gaussian, to become bimodal (Figure S1B). Plasticity also influences the distribution of oscillator units, resulting in bimodal distributions of both phase and phase difference (Figure S1C), consistent with previous studies that incorporated the Hebbian rule into the Kuramoto model^21,26^. This bimodal distribution of the coupling weight deviates from the long-tailed skewed distributions typically observed in biological synaptic weights^10,32,33^.

To address this discrepancy, we introduced an exponential-based Hebbian plasticity rule to adjust the decay rate (sparseness parameter *γ*) of the gain for the phase difference (Figure 1D). We applied this rule to a range of parameter sets and evaluated the resulting *J*_*ee*_ distribution using skewness, kurtosis, and dip values, which quantify the distribution bimodality^34^. Although *J*_*ee*_ distributions initially followed a Gaussian pattern, they transitioned into various bimodal or skewed forms (Figure 1E and F). We also quantified the synchronization of the excUnits using the mean and standard deviation of the order parameter *R*_*exc*_ over time, denoted as E[*R*_*exc*_]_t_ and S[*R*_*exc*_]_t_, respectively. Plasticity in *J*_*ee*_ affects synchronization (Figure 1G), resulting in uniform, bimodal, or unimodal phase and phase-difference distributions depending on the plasticity parameters (Figure S1D). Moreover, the boundary of the statistical features of the *J*_*ee*_ distribution (Figure 1E) appears to coincide with the synchronization boundary (Figure 1G). For further analysis, we chose a parameter set that resulted in a long-tailed skewed *J*_*ee*_ distribution (Figure 1F, top, magenta circle), which more closely reflects the synaptic weight distributions observed in the brain^10,32,33^.

### Coupling fluctuations increase in the bistable state

We next investigated how distinct dynamic states associated with different EI balances affect coupling weights. To this end, we altered the EI balance of the pEI-Kuramoto model by manipulating the interaction strength from inhUnits. We unified *K*_*ie*_ and *K*_*ii*_ into a single parameter, *K*_*i*_, to simplify the simulation, following the approach used in our previous study^14^. We then examined how the *J*_*ee*_ fluctuations were associated with the *K*_*i*_ values, quantified as the standard deviation of *J*_*ee*_ over time (S[*J*_*ee*_]_t_; see Materials and Methods) (Figure 2A and B). We determined the system’s dynamic state by analyzing the time course of *R*_*exc*_ (Figure 2C and D), as previously described^14^.

**Figure 2.**
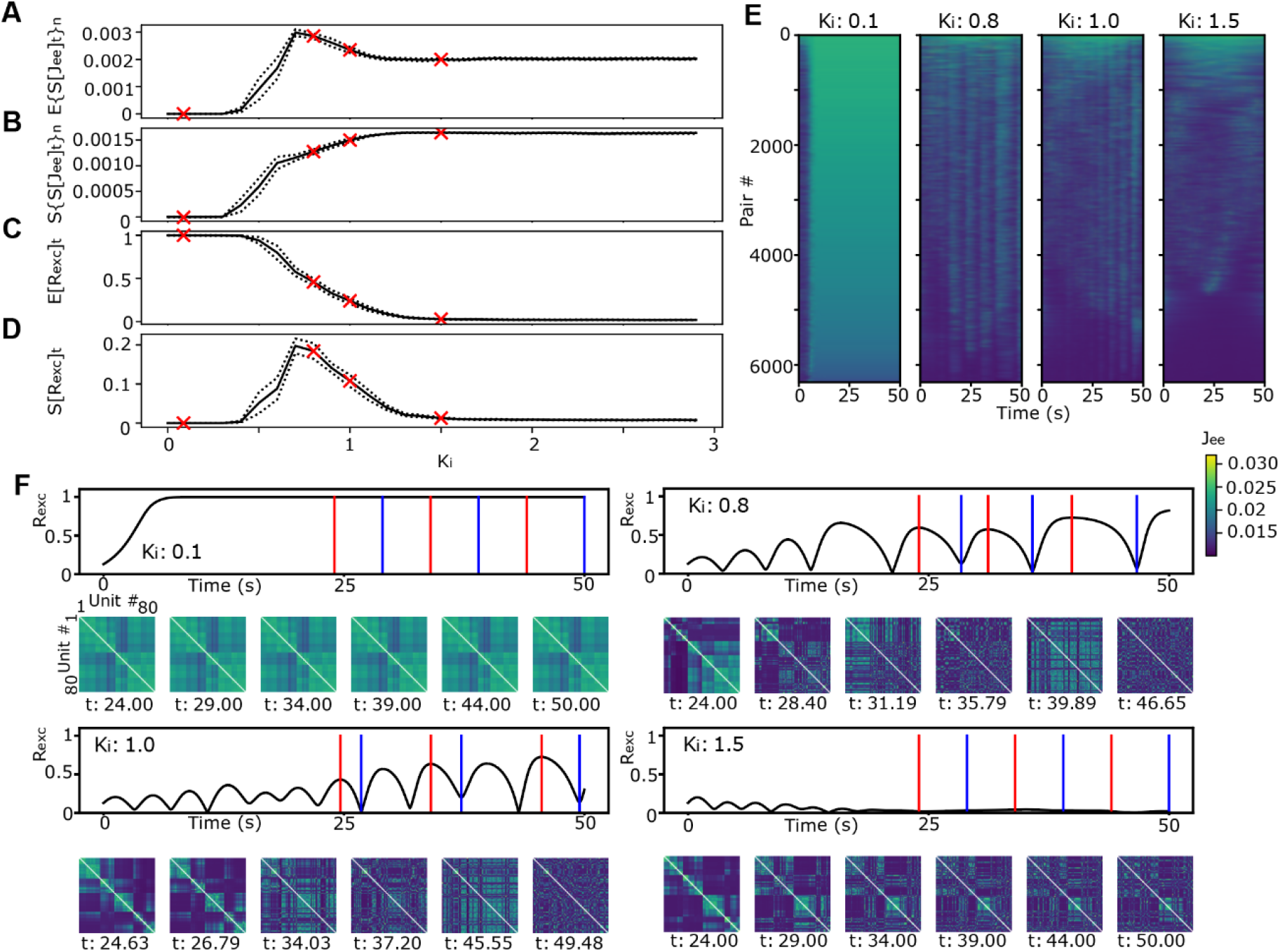
Changes in *J*_*ee*_ fluctuation with *K*_*i*_ modification. (A–D) (A) E{S[*J*_*ee*_]_t_}_n_, (B) S{S[*J*_*ee*_]_t_}_n_, (C) E[*R*_*exc*_]_t_, and (D) S[*R*_*exc*_]_t_ as a function of *K*_*i*_. Red crosses indicate each value in the *K*_*i*_ conditions shown in (E) and (F). (E) Time course of *J*_*ee*_ weights under different *K*_*i*_ conditions. The order of unit pairs is sorted by E[*J*_*ee*_]_t_ for each *K*_*i*_ value. (F) Upper panels show *R*_*exc*_ traces. The lower panels show the *J*_*ee*_ weight matrix at selected time points indicated by the red and blue vertical lines in the top panels (red and blue are local maximum and local minimum, respectively, for *K*_*i*_ = 0.8 and 1.0; *t* = 24, 29, 34, 39, 44, and 50 for *K*_*i*_ = 0.1 and 1.5). The order of units is rearranged by the result of hierarchical clustering for the leftmost panel (around *t*: 24-25). E and F share the color bar. *μ*_*Jee*_ = 1, *σ*_*Jee*_ = 0.1, *K*_*ei*_ = 3, *α* = 0.01, *β* = 0.5, *γ* = 2.

When *K*_*i*_ < 0.4, both the average (E{S[*J*_*ee*_]_t_}_n_) and standard deviation (S{S[*J*_*ee*_]_t_}_n_) of S[*J*_*ee*_]_t_ across unit pairs were almost zero (Figure 2A and B), indicating no plastic changes in *J*_*ee*_. In this *K*_*i*_ range, the system was in a synchronized state, characterized by high E[*R*_*exc*_]_t_ and low S[*R*_*exc*_]_t_ (Figure 2C and D). The coupling weights between individual pairs exhibited no fluctuation over time after reaching the synchronized state (Figure 2E and F, *K*_*i*_: 0.1; Video S1).

As *K*_*i*_ increased, E{S[*J*_*ee*_]_t_}_n_ also increased, peaking at *K*_*i*_ = 0.7, then decreased, and eventually stabilized around *K*_*i*_ = 1.5 (Figure 2A). Meanwhile, the S{S[*J*_*ee*_]_t_}_n_ continued to increase (Figure 2B). For *K*_*i*_ values above 1.5, the system transitioned into a desynchronized state characterized by low E[*R*_*exc*_]_t_ and S[*R*_*exc*_]_t_ values (Figure 2C and D). Strong or weak couplings remained stable over time, whereas those of intermediate strength fluctuated (Figure 2E, *K*_*i*_: 1.5; Video S2). The system generated transient clusters of units coupled with strong *J*_*ee*_ weights (Figure 2F, *K*_*i*_: 1.5, the lower leftmost panel, *t* = 24). Over time, most units left these clusters, leaving only small, stable subsets (Figure 2F, *K*_*i*_: 1.5, the lower panels, *t* = 29–50).

In the intermediate range (0.4 < *K*_*i*_ < 1.5), the system exhibited a bistable state with high S[*R*_*exc*_]_t_ (Figure 2D). Notably, the S[*J*_*ee*_]_t_ was significantly higher in this bistable state (Figure 2A). Similar to the desynchronized state (*K*_*i*_ > 1.5), the couplings with high or low weights were stable. However, the intermediate-strength couplings fluctuated more (Figure 2E and F, *K*_*i*_: 0.8 and 1.0). Notably, the number of these intermediate fluctuating couplings was higher in the intermediate *K*_*i*_ range (Figure 2E and F, *K*_*i*_: 0.8 and 1.0; Videos S3 and S4) than in the low and high *K*_*i*_ ranges. Thus, transitions in the dynamic state were accompanied by changes in the coupling weight fluctuations. Particularly, the bistable state tended to exhibit larger *J*_*ee*_ fluctuations.

Note that Hebbian plasticity affected the dynamic state of the pEI-Kuramoto model (Figure S2A), slightly shifting the bifurcation points between the synchronized and bistable states, and the bistable and desynchronized states toward larger *K*_*i*_ values. Owing to this effect, we observed a small discrepancy from the state predicted by the eigenvalue-based classification for the EI-Kuramoto model^14^ (data not shown). However, the overall trend of the state transition from synchronized to bistable and then to desynchronized as *K*_*i*_ values increased remained consistent across different Hebbian learning rates. The enhancement of synchronization induced by Hebbian plasticity was consistent with the findings of previous reports on Kuramoto models incorporating cosine-shaped Hebbian plasticity^26,31^. In contrast, *σ*_*Jee*_ did not affect the dynamic states (Figure S2B), consistent with the results in the absence of *J*_*ee*_ plasticity shown in Figure 1C.

### Intermediate couplings are selectively destabilized while the strong couplings remain stable

We further investigated coupling weight fluctuations under conditions of slightly weaker inhibition than in the desynchronized state, assuming intermediate brain states between wake-like and sleep/rest-like states. We analyzed the individual *J*_*ee*_ coupling weights at selected points of *K*_*i*_ (*K*_*i*_ = 0.8, 1.0, and 1.5, red crosses in Figure 2A–D, except *K*_*i*_ = 0.1). Histograms of E[*J*_*ee*_]_t_ (mean of *J*_*ee*_ over time for each pair of units) and S[*J*_*ee*_]_t_ also tended to shift toward higher values as *K*_*i*_ decreased (Figure 3A and B). A scatter plot of E[*J*_*ee*_]_t_ and S[*J*_*ee*_]_t_ formed a characteristic “bow-shaped” distribution (Figure 3C). In this two-dimensional distribution, data points near the minimum and maximum E[*J*_*ee*_]_t_ values had lower S[*J*_*ee*_]_t_ values, whereas intermediate E[*J*_*ee*_]_t_ showed a higher S[*J*_*ee*_]_t_. This pattern was consistent across all distributions for different *K*_*i*_ values. A similar phenomenon, the stabilization of strong couplings (e.g., synapses with large mushroom-shaped spines), has been observed in brain synapses^35–37^.

**Figure 3.**
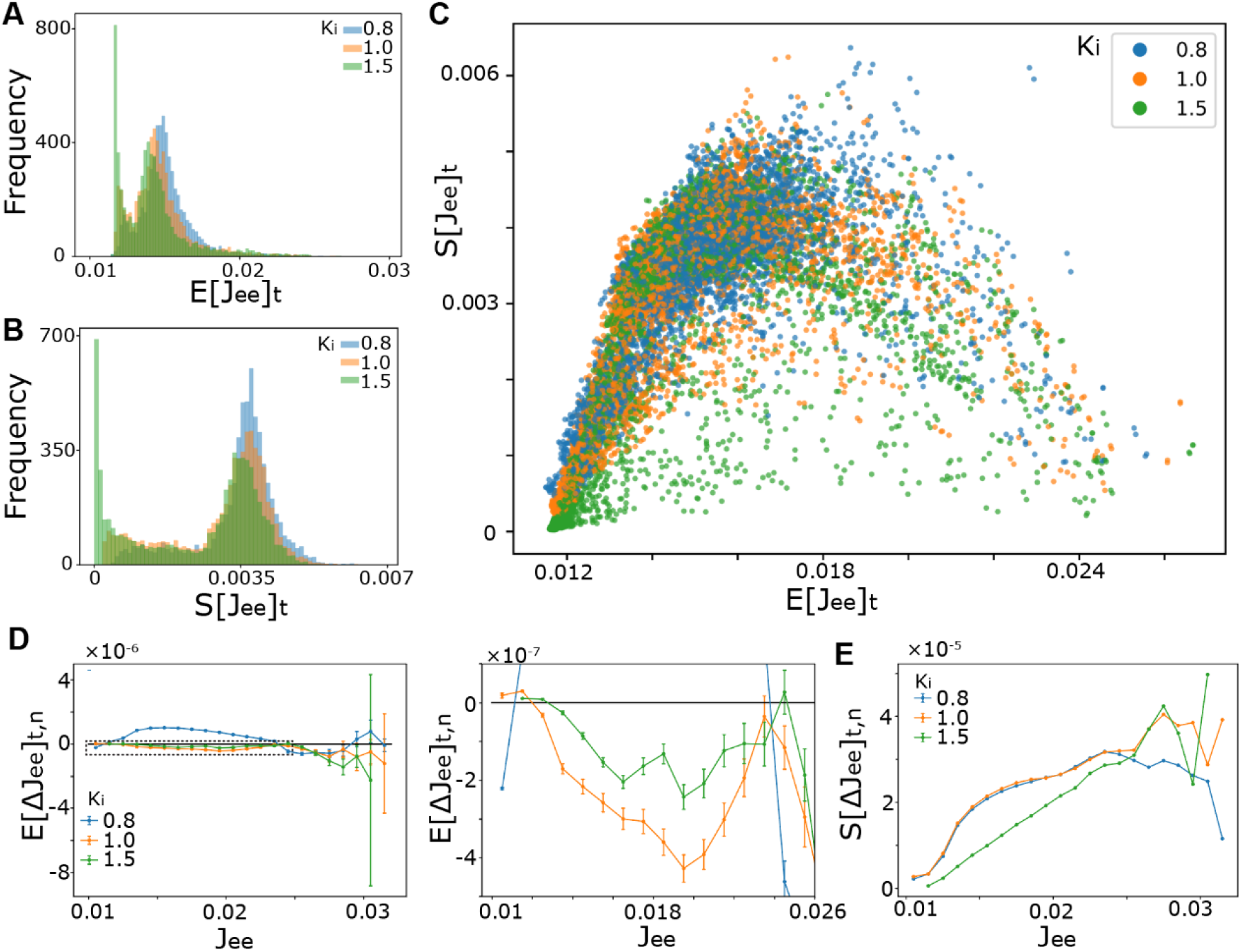
Relationship between the magnitude and fluctuation of *J*_*ee*_. (A) E[*J*_*ee*_]_t_ histogram. (B) S[*J*_*ee*_]_t_ histogram. (C) E[*J*_*ee*_]_t_ vs. S[*J*_*ee*_]_t_ scatter plot for different *K*_*i*_ values. (D) E[*ΔJ*_*ee*_]_t,n_ as a function of *J*_*ee*_ for different *K*_*i*_ values. The right panel shows an enlarged view of the region enclosed by the dotted rectangle in the left panel. Black line indicates E[*ΔJ*_*ee*_]_t,n_ = 0. Error bars indicate the standard error of the mean (SEM). (E) S[*ΔJ*_*ee*_]_t,n_ as a function of *J*_*ee*_ for different *K*_*i*_ values. At each time point, all pairs of *ΔJ*_*ee*_ and the immediately preceding *J*_*ee*_ values were collected, binned by the *J*_*ee*_ values, and used to compute the statistics within each bin (D, E). *μ*_*Jee*_ = 1, *σ*_*Jee*_ = 0.1, *K*_*ei*_ = 3, *α* = 0.01, *β* = 0.5, *γ* = 2.

Furthermore, the S[*J*_*ee*_]_t_ values tended to increase at intermediate E[*J*_*ee*_]_t_ values with decreasing *K*_*i*_ (Figure 3C), suggesting that *K*_*i*_ can regulate the fluctuations in individual *J*_*ee*_ plastic couplings. At high *K*_*i*_ values in desynchronized states (*K*_*i*_ = 1.5, green dots), the couplings were relatively stable, whereas at low *K*_*i*_ values in bistable states (*K*_*i*_ = 1.0 and 0.8, orange and blue dots), the couplings of intermediate weights were more unstable. Strong *J*_*ee*_ coupling remained stable with small fluctuations, even when *K*_*i*_ was low. These dynamics may provide a mechanism that preserves the coupling of particularly strong pairs and resets moderate-weight pairs, facilitating the reorganization of the network structure (Figure 2E and 2F).

To enable a direct comparison with Figure 7A and 7B in Yasumatsu et al^37^, which report the relationship between spine volume and changes in spine volume in hippocampal slices, we quantified the change in coupling weight, *ΔJ*_*ee*_ (= *J*_*ee*_(*t+Δt*) – *J*_*ee*_(*t*)), as a function of its current value, *J*_*ee*_, at each time step in our pEI-Kuramoto model (Figure 3D and E). We pooled *J*_*ee*_ values and their changes *ΔJ*_*ee*_ across all excUnit pairs and time points, binned the *J*_*ee*_ values, and computed the mean and the standard deviation of *ΔJ*_*ee*_ for each *J*_*ee*_ bin (E[*ΔJ*_*ee*_]_t,n_ and S[*ΔJ*_*ee*_]_t,n_). In the desynchronized state (*K*_*i*_ = 1.5, green lines), E[ΔJ_ee_]_t,n_ was slightly but significantly positive for small *J*_*ee*_ (Wilcoxon signed-rank test, bin=[0.011,0.012), mean=1.1×10^-8^, n=6,686,034, W=1.06×10^13^, p<1.0×10^-308^), then decreased with increasing *J*_*ee*_, reaching a local minimum near *J*_*ee*_≈0.02. Subsequently, E[*ΔJ*_*ee*_]_t,n_ increased and approached zero around *J*_*ee*_≈0.025 (Figure 3D), a region corresponding to the peak of E[*J*_*ee*_]_t_ (Figure 3A and C). S[*ΔJ*_*ee*_]_t,n_ increased monotonically over the same range of *J*_*ee*_ (Figure 3E). These trends closely mirror observations in hippocampal slices and *in vivo* cortical imaging^37,38^. In comparison, the bistable regimes (*K*_*i*_=0.8 and 1.0, orange and blue lines) showed large deviations in the mean change (E[Δ*J*_*ee*_]_t,n_) either in the same direction (*K*_*i*_=1.0, orange line) or the opposite direction (*K*_*i*_=0.8, blue line) as the desynchronized state (Figure 3D). Furthermore, the standard deviation (S[*ΔJ*_*ee*_]_t,n_) was generally elevated relative to the desynchronized state across the range of *J*_*ee*_ from 0.01 to 0.025 (Figure 3E).

### Factors determining coupling weight properties

To identify the parameters that determine the individual *J*_*ee*_ coupling properties (E[*J*_*ee*_]_t_ and S[*J*_*ee*_]_t_) shown in Figure 3, we explored the candidate parameters (natural frequency difference *Δω*, initial phase difference *Δθ*, and initial *J*_*ee*_ values of each excUnit pair) that may contribute to *J*_*ee*_ fluctuations (Figure 4). We colored the dots of E[*J*_*ee*_]_t_ vs. S[*J*_*ee*_]_t_ distribution (Figure 3C) according to each parameter value (Figure 4A–C). For statistical comparison, couplings were roughly categorized based on whether E[*J*_*ee*_]_t_ exceeded 0.018 (high-E[*J*_*ee*_]_t_ group) or not (low-E[*J*_*ee*_]_t_ group), because the correlation between E[*J*_*ee*_]_t_ and S[*J*_*ee*_]_t_ reversed around this point (Figure 4D–F).

**Figure 4.**
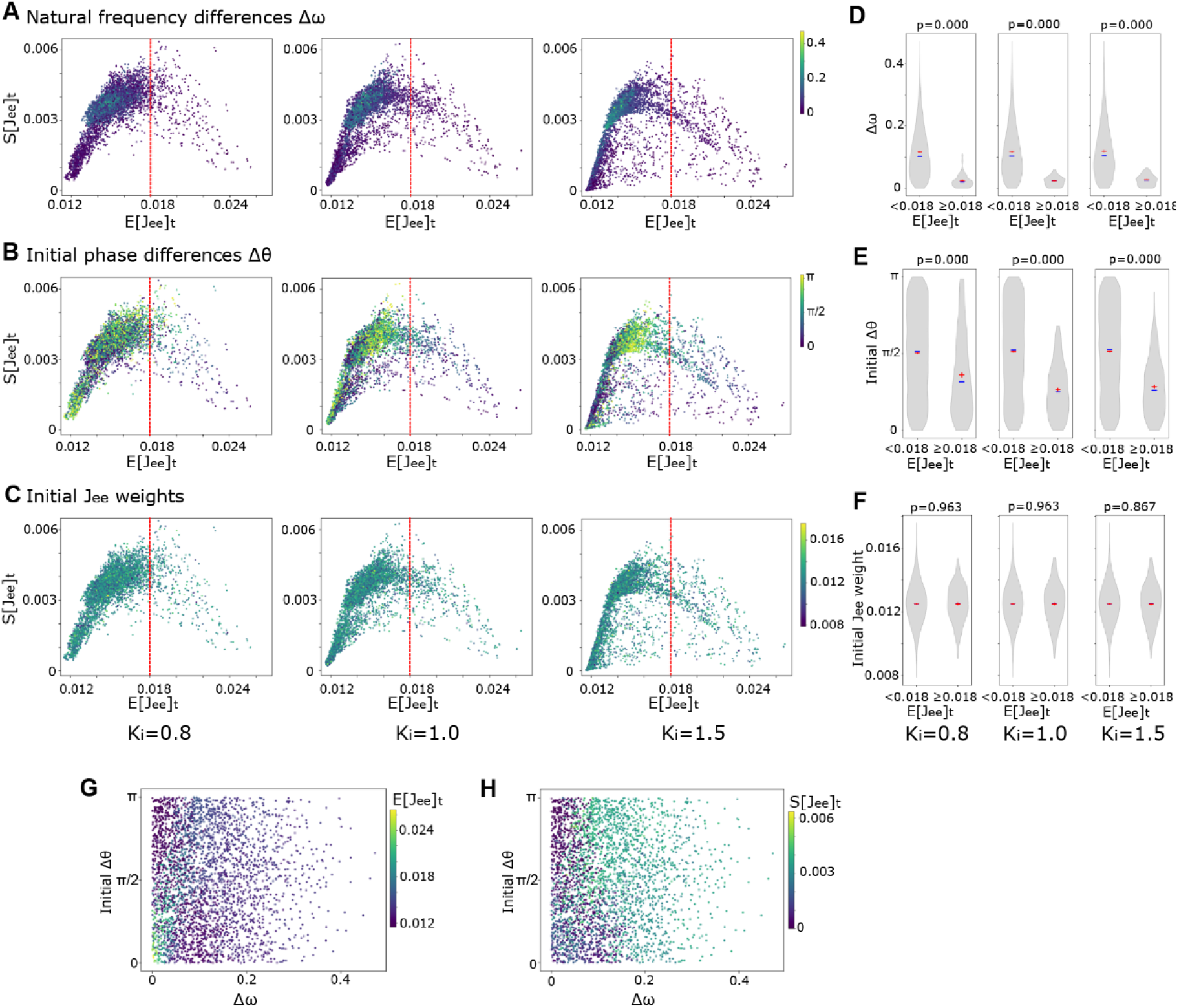
Factors affecting the *J*_*ee*_ fluctuation. (A–C) E[*J*_*ee*_]_t_ vs. S[*J*_*ee*_]_t_ scatter plot with color coding of (A) the natural frequency difference *Δω*, (B) the initial phase difference *Δθ*, and (C) the initial *J*_*ee*_ weight over different *K*_*i*_. The red dotted line for each panel indicates E[*J*_*ee*_]_t_ = 0.018. Left column, *K*_*i*_ = 0.8; middle column, *K*_*i*_ = 1.0; right column, *K*_*i*_ = 1.5. (D–F) Violin plots provide distribution densities based on whether the threshold E[*J*_*ee*_]_t_ ≥ 0.18 or not for (D) natural frequency differences *Δω*, (E) initial phase differences *Δθ*, and (F) initial *J*_*ee*_ weights across different *K*_*i*_. The red and blue horizontal lines indicate the mean and median values, respectively. The red vertical lines indicate the standard errors. The numbers in the upper right corner of each panel are the p-values from the Mann–Whitney U-test. The numbers of unit pairs were as follows: for *K*_*i*_ = 0.8, E[*J*_*ee*_]_t_ < 0.018: *n* = 5,968, E[*J*_*ee*_]_t_ ≥ 0.018: *n* = 352; for *K*_*i*_ = 1.0, E[*J*_*ee*_]_t_ < 0.018: *n* = 5,920, E[*J*_*ee*_]_t_ ≥ 0.018: *n* = 400; for *K*_*i*_ = 1.5, E[*J*_*ee*_]_t_ < 0.018: *n* = 5,852, E[*J*_*ee*_]_t_ ≥ 0.018: *n* = 468. The statistics were as follows: (D, natural frequency difference), *K*_*i*_ = 0.8, *z* = 24.7, *p* < 0.001; *K*_*i*_ = 1.0, *z* = 26.8, *p* < 0.001; *K*_*i*_ = 1.5, *z* = 27.8, *p* < 0.001. (E, initial phase difference) *K*_*i*_ = 0.8, *z* = 9.20, *p* < 0.001; *K*_*i*_ = 1.0, *z* = 16.4, *p* < 0.001; *K*_*i*_ = 1.5, *z* = 16.7, *p* < 0.001. (F, initial *J*_*ee*_ weight) *K*_*i*_ = 0.8, *z* = –0.046, *p* = 0.96; *K*_*i*_ = 1.0, *z* = 0.046, *p* = 0.96; *K*_*i*_ = 1.5, *z* = 0.17, *p* = 0.87. (G and H) Scatter plots of natural frequency differences *Δω* and initial phase differences *Δθ* with color coding of (G) E[*J*_*ee*_]_t_ and (H) S[*J*_*ee*_]_t_ (*K*_*i*_ = 1.5). *μ*_*Jee*_ = 1, *σ*_*Jee*_ = 0.1, *K*_*ei*_ = 3, *α* = 0.01, *β* = 0.5, *γ* = 2.

Regarding differences in the natural frequency *Δω*, most pairs in the high (≥0.018) E[*J*_*ee*_]_t_ group had relatively small *Δω*, whereas pairs with larger *Δω* tended to be in the low (<0.018) E[*J*_*ee*_]_t_ group (Figure 4A and 4D). In the low-E[*J*_*ee*_]_t_ group, S[*J*_*ee*_]_t_ seemingly increased with *Δω* (Figure 4A). The initial phase differences *Δθ* between the pairs were also significantly different between the high- and low-E [*J*_*ee*_]_t_ groups (Figure 4B and E). In the high-E[*J*_*ee*_]_t_ group, most pairs exhibited small initial *Δθ*, whereas in the low-E[*J*_*ee*_]_t_ group, the initial *Δθ* was uniformly distributed. Additionally, we examined the effect of the initial *J*_*ee*_ coupling weight, which showed no noticeable bias (Figure 4C) and followed a normal distribution in both the high and low-E[*J*_*ee*_]_t_ groups (Figure 4F). Considering these results, pairs with small differences in natural frequency and initial phase positions maintained more stable and larger *J*_*ee*_ weights (Figure 4G and H). Conversely, the initial *J*_*ee*_ weight was not an important factor in determining the stability and strength of the couplings.

### Theoretical analysis of quasi-steady coupling weight distribution

We investigated the theoretical mechanism underlying the fluctuation of coupling weights *J*_*ee*_ by analyzing their distribution in a quasi-steady state, assuming *J*_*ee,kj*_(*t* + Δ*t*) ≈ *J*_*ee,kj*_(*t*) (Supplemental Text). Although a true static steady weight distribution does not exist in the desynchronized or bistable states due to the non-constant phases of each oscillator unit, the desynchronized condition allows for a quasi-steady state where the phase and weight distributions evolve on a sufficiently slow timescale, resulting in minimal fluctuation of the *J*_*ee*_ distribution. Under this assumption, the coupling weights do not change as a result of the plastic modulation in Equations M11 and M14, leading to the following relationship (Supplemental Text, Equation S3).

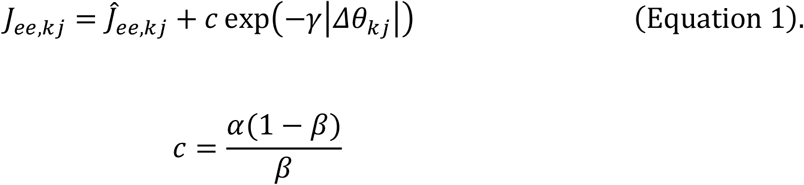

The first term, 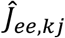, represents the value of *J*_*ee,kj*_, which is rescaled with the homeostatic regulation plasticity rule so that the mean and variance become 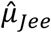 and 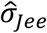, respectively. The second term is the Hebbian potentiation component, scaled by the ratio between the Hebbian and homeostatic plasticity learning rates *c*. This equation indicates that the weight *J*_*ee,kj*_ depends on the distribution of 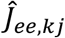 and the distribution of phase difference between unit *k* and *j Δθ*_*k*j_ (*c* exp(−*γ*|Δ*θ*_*kj*_|)). Assuming that this rule remains applicable during the approach to a quasi-steady state, successive iterations of the update rule progressively integrate the second *Δθ*-dependent Hebbian term into the *J*_*ee*_ distribution. Furthermore, the distribution of *Δθ* is dependent on that of *θ* (Supplemental Text). Therefore, when the weight dynamics are sufficiently slow, the *J*_*ee*_ distribution is determined by the phase distribution. Indeed, the characteristics of the *J*_*ee*_ distribution appear to be correlated with the synchronization state of the system, which reflects the shape of the phase distribution (Figure 1E, 1F, and S1). Specifically, a uniform phase (and phase difference) distribution (desynchronized state) corresponds to a right-skewed, long-tailed *J*_*ee*_ distribution (magenta circle in Figure 1E, F and S1D). A bimodal phase distribution corresponds to a bimodal *J*_*ee*_ distribution (green circle in Figure 1E, F and S1D and the cos rule in S1B and S1C). A unimodal phase distribution (synchronized state) corresponds to a left-skewed *J*_*ee*_ distribution (yellow circle in Figure 1E, F and S1D).

Next, we considered the change in the phase difference for a pair of excUnits, *Δθ*_*kj*_. Assuming that *J*_*ee,kj*_ and *Δθ*_*kj*_ are constant, the following relationship can be derived from Equations 1 and M9 (Supplemental Text, Equation S18):

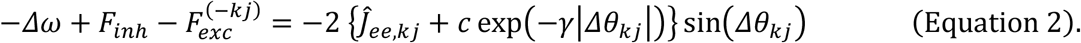

*Δω* is the natural frequency difference between units *k* and *j, F*_*inh*_ is the interaction strength from the inhUnits, and 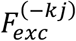 is the interaction strength from the excUnits, except for units *k* and *j*. 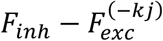 is the *R*_*exc*_- and *R*_*inh*_-dependent term; its direction can be either attractive or repulsive, depending on the phase configuration of the system (Supplemental Text). The right-hand side of this equation represents the interaction strength between units *k* and *j*, whereas the left-hand side represents the net forces of the other factors. A stable fixed point for *Δθ*_*kj*_ exists, where the functions for the left- and right-hand sides intersect (Figure S3).

On the right-hand side, the attractive force is driven by both the homeostatic set-point, 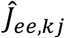, and the Hebbian potentiation term. Near *Δθ*≈0, the Hebbian effect is strong, yielding a large attractive force. As the phase difference ∣*Δθ*∣ increases, this Hebbian potentiation weakens, and the coupling weight relative position within the *J*_*ee*_ distribution falls to a lower value, causing an overall attractive force reduction.

On the left-hand side, while the influence of *F*_*inh*_ and 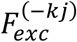 is limited in a desynchronized state where *R*_*exc*_ and *R*_*inh*_ are near zero, their impact becomes non-negligible in the bistable state, where these order parameters undergo periodic modulation. Sufficiently large fluctuations in these forces cause the stable fixed point to disappear or be substantially displaced, thereby leading to large fluctuations in the phase difference between the unit pair.

### Cyclic modulation of inhibition

Finally, we varied *K*_*i*_ to mimic the sleep–wake cycle^39,40^. *J*_*ee*_ fluctuated (Figure 5A, Video S5) in response to *K*_*i*_ modulation (Figure 5B, top panel). *R*_*exc*_ alternated between the desynchronized and bistable states (Figure 5B, middle), with corresponding changes in E{S[*J*_*ee*_]_t_}_n_ (Figure 5B, bottom). When *K*_*i*_ was high, the system was in a desynchronized state, and E{S[*J*_*ee*_]_t_}_n_ was small. Conversely, when *K*_*i*_ was low, the system exhibited a bistable state, and E{S[*J*_*ee*_]_t_}_n_ increased. These dynamics can be considered as modeling how the circuit structure is stabilized during the wake-like state and destabilized during the sleep/rest-like state, thereby promoting circuit reorganization.

**Figure 5.**
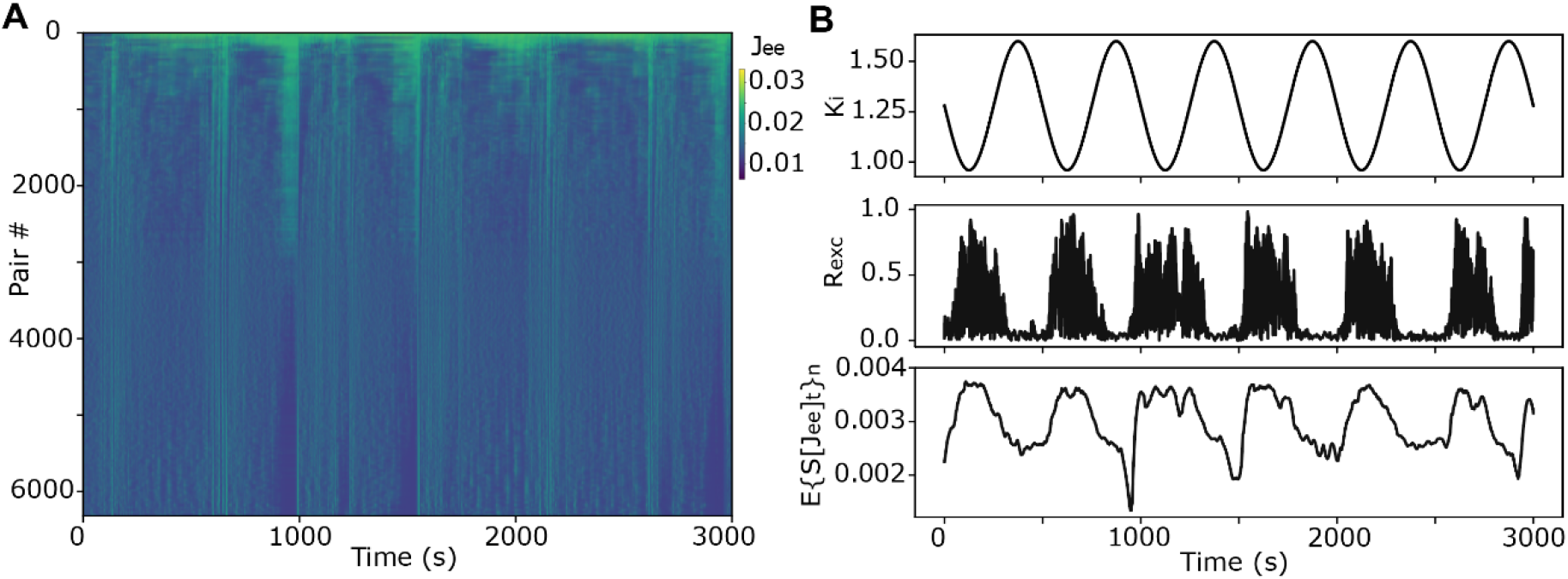
*J*_*ee*_ fluctuation changes with *K*_*i*_ cyclic modulation. (A) Time trace of each *J*_*ee*_ weight sorted by E[*J*_*ee*_]_t._ (B) Time traces of (top) *K*_*i*_ modulation, (middle) *R*_*exc*_, and (low) E{S[*J*_*ee*_]_t_}_n_. *μ*_*Jee*_ = 1, *σ*_*Jee*_ = 0.1, *K*_*ei*_ = 3, *α* = 0.01, *β* = 0.5, *γ* = 2.

## Discussion

The main conceptual advance of this study is to show that EI balance can determine not only the synchronization state of a network but also whether plastic couplings are stabilized or rendered labile. In the model, strong inhibition favored a desynchronized regime with relatively stable couplings, whereas weaker inhibition produced a bistable regime that selectively destabilized intermediate couplings while preserving the strongest ones. This provides a candidate systems-level mechanism by which sleep/rest-like network states can promote reorganization without erasing established strong connections.

Although the sleep/rest-like state is often associated with synchrony, it is not uniformly synchronized. Near the transition between synchronized and desynchronized states, the dynamics tend to become irregular or weakly stable, as captured by two-population coupled-oscillator models ^41,42^. Sleep onset can be formalized as a noisy saddle-node bifurcation ^43^, and macroscopic brain-state transitions are often argued to occur near the bifurcation point ^44,45^. Beyond the transitions between wakefulness and sleep, even within sleep, the microarchitecture exhibits intrinsic instability, such as the NREM-REM sleep cycle^3,4,39,40^, the cyclic alternating pattern in NREM sleep^46^, UP–DOWN state alternations^47,48^, and microarousals^49^, which may reflect switching among synchronized and desynchronized states. These fluctuation-driven transitions align with the bistable regime of our model under reduced inhibition.

Preserving robust couplings while randomizing weaker ones simplifies the network, rendering it sparse. The plasticity parameters used here, which yield a long-tailed coupling-weight distribution (Figure 1E and F) similar to that observed in the brain^10,32,33^ and mirror the EI balance during state transitions and in sleep/rest-like states^7–10^, appear to facilitate network sparsification. The selective preservation of strong synaptic connections is also frequently observed in the brain^35–37^ and is understood through molecular mechanisms, including the dynamics of NMDA receptors^5,50^ and actin filaments^51^. Our results introduce a complementary, system-level mechanism for sustaining strong coupling, viewed through the dynamics of collective oscillators. Additionally, our results may be mechanistically linked to the systems-level observation that only a subset of neuronal populations remain stable while others drift^52–54^. This process resembles pruning strategies used in artificial neural networks that facilitate knowledge generalization^55,56^. Such pruning-like dynamics are particularly relevant during sleep/rest-like states when synaptic downregulation is dominant^5,50^, as fluctuations in coupling strength can weaken intermediate connections and eventually reset them toward baseline.

Fluctuations in network connectivity can promote network reorganization. In our model, bistable states under low inhibition conditions, mirroring the EI balance during state transitions and sleep/rest-like states^7–10^, promoted fluctuating coupling weights between excUnits. It has also been proposed that the brain enhances neuronal circuit reorganization during the sleep/rest-like state^3,4^. Combined with mechanisms that evaluate and stabilize reorganized circuits^57^—potentially including dopaminergic signaling during NREM sleep^58^, which may provide a reinforcement-like signal and gate plastic changes—the sleep/rest-like state could, in principle, contribute to the generation of novel associations, creative ideas, or innovative problem-solving strategies.

The characteristic “bow-shaped” relationship between coupling weight mean (E[*J*_*ee*_]_t_) and standard deviation (S[*J*_*ee*_]_t_) (Figure 3C) arises from the stability of phase differences between excUnits (Equation 2). Pairs with strong coupling maintain small and stable phase differences (*Δθ*≈0), ensuring that the Hebbian potentiation term (c exp(−γ|Δθ|)) and their relative position in the *J*_*ee*_ distribution remain consistently high. In contrast, pairs with weak couplings evolve independently with fluctuations driven mainly by natural frequency differences *Δω*. The largest S[*J*_*ee*_]_t_ was found in pairs with intermediate weights. In this regime, the intrinsic synchronizing force between the two units is closely counterbalanced by external driving influences, chiefly their natural frequency difference, *Δω*. Furthermore, Hebbian potentiation and homeostatic depression are also balanced. Small fluctuations in *Δθ* can frequently tip these balances and reverse the direction of the weight change, leading to large fluctuations in the coupling weight *J*_*ee*_.

From a dynamics perspective, these mechanisms explain not only the characteristic bow-shaped distribution in Figure 3C but also a key feature of biological synapses: larger synapses are more stable than smaller ones^35–37^. Moreover, particularly in the desynchronized state, our model reproduces the relationship between synaptic size and changes in synaptic size reported in experimental studies^37,38^ (Figure 3D and E, green lines; see Results), further supporting its biological relevance.

The quasi-steady *J*_*ee*_ distribution is determined by the phase-difference distribution (Figure 1E, 1F, S1C and S1D), as it reflects a transformation of the *Δθ* distribution via the Hebbian function (Equation 1). In this study, the exponential rule *c* exp(−*γ*|Δ*θ*_*kj*_|) transforms the uniform phase-difference distribution into a hyperbolic weight distribution (Supplemental Text, Equation S8–12). Modifying the Hebbian function would allow the generation of alternative *J*_*ee*_ distributions, such as log-normal distributions commonly observed in the brain^10,32,33^.

Equation 2 further predicts that unit pairs with small natural frequency differences (*Δω*) and small initial phase differences (*Δθ*) are more robust and more likely to return to a stable, synchronized state even after large perturbations, consistent with the results in Figure 4. In our simulation (Figure 1F, magenta circle), the Hebbian plasticity-induced enhancement of interaction strength near *Δθ*=0 is limited, even though the plasticity slightly strengthens the restoring force near the stable point (Figure S3). Thus, the tendency for pairs with small *Δω* and small initial *Δθ* to be more robust and stable arises primarily from the underlying sinusoidal form of the baseline interaction function rather than plasticity-induced modifications. This finding resonates with experimental observations that neurons exhibiting synchronized firing prior to learning (analogous to a small *Δθ*) are preferentially incorporated into the same assembly during learning^59–61^.

Fluctuations in *J*_*ee*_ at intermediate *K*_*i*_ levels (Figs. 2–4, *K*_*i*_=0.8, 1.0) arise from bistability between synchronized and desynchronized states. This alternation perturbs phase distributions, which directly drive changes in *J*_*ee*_. Concurrent periodic fluctuations of the order parameters *R*_*exc*_ and *R*_*inh*_ constantly modulate the interaction term, 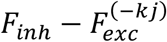 (Equation 2), further destabilizing phase differences *Δθ* of moderately coupled pairs.

While the Kuramoto model provides conceptual insight, it is substantially simpler than biological neural circuits. The model reduces neuronal interactions to attractive and repulsive forces, neglecting pulse-based communication, conduction delays, and complex network topology^62^. Additionally, our model simplifies the complexity of neuroplasticity—excluding neurotransmitters^63^, inhibitory plasticity^64,65^, and adaptation^66^—and does not distinguish between specific sleep stages (e.g., NREM vs. REM), which are thought to play distinct roles in memory processing and circuit reorganization^3,4^. Addressing these limitations is essential for a comprehensive understanding of neural dynamics.

Future work can extend the framework by introducing explicit evaluation and stabilization signals that select among reorganized circuits^57,67^, potentially linking fluctuation-driven reorganization to selective retention during sleep and rest. Another important direction is to examine how synchrony transitions influence interregional plasticity between functionally distinct networks, such as hippocampus and cortex^68^, thereby linking circuit-level reorganization to systems-level consolidation.

## Materials and Methods

### pEI-Kuramoto model

The original Kuramoto model^11,12,69^ is expressed as:

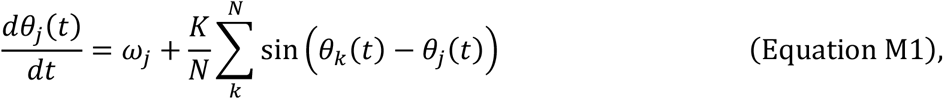

where *θ*_*j*_(*t*) is the phase of unit *j* at time *t, ω*_*j*_ is the natural frequency of unit j following a probability distribution *g*(*ω*), *N* is the number of units, and *K* is the interaction strength among the oscillation units. In the original Kuramoto model, *K* is applied constantly and uniformly to all unit pairs. The order parameter *R* and the mean phase *ψ* are then calculated from the mean vector of the oscillators as:

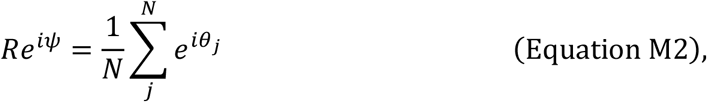

where *i* denotes an imaginary unit. Given Euler’s formula and Equation M2,

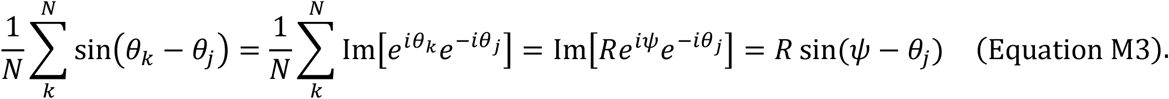

Considering Equation M3, Equation M1 becomes

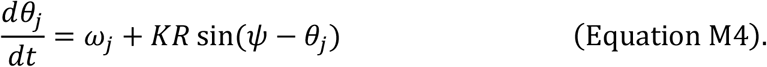

In our previous EI-Kuramoto model^14^, the oscillation units were categorized into two groups: excitatory and inhibitory units, referred to as excUnits and inhUnits, respectively. Employing these terms clarified that the units in the model represented excitatory and inhibitory factors, distinguishing them from actual neurons or synapses. Zero-phase-lag sine interactions are purely attractive for excUnits and purely repulsive for inhUnits. Hence, the terms “excitatory” and “inhibitory” are used interchangeably with “attractive” and “repulsive,” respectively. The interaction strength *K* is divided into four quantifiable parameters: *K*_*ee*_ (attractive strength between excUnits), *K*_*ii*_ (repulsive strength between inhUnits), *K*_*ei*_ (attractive strength from excUnits to inhUnits), and *K*_*ie*_ (repulsive strength from inhUnits to excUnits). *K*_*ee*_, *K*_*ii*_, *K*_*ei*_, and *K*_*ie*_ are distinct positive real numbers that are uniformly applicable across all the unit pairs. Consequently, Equation M4 becomes:

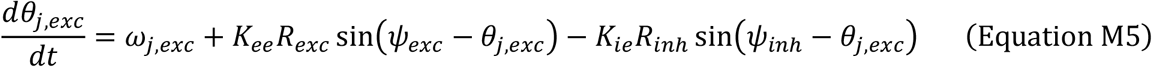

for excUnits, and

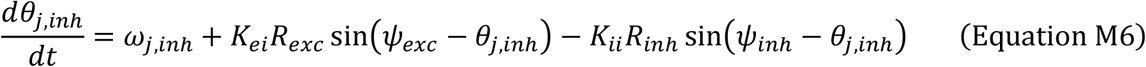

for inhUnits, where exc and inh represent the parameters for excUnits and inhUnits, respectively. The order parameter and mean phase of each excUnit and inhUnit are

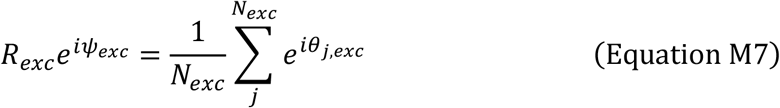

and

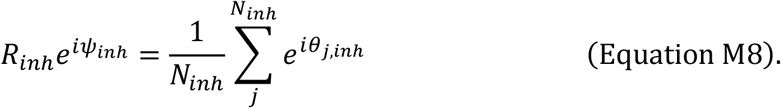

In the present study, we developed a pEI-Kuramoto model (Figure 1A). A key modification of our previous EI-Kuramoto model is that the constant and uniform interaction strength value *K*_*ee*_ is replaced with the heterogeneous and plastic coupling weight (interaction strength of an individual unit pair) matrix *J*_*ee*_. The *J*_*ee*_ distribution is defined by its mean *μ*_*Jee*_ and standard deviation *σ*_*Jee*_, which are scaled with the inverse of *N*_*exc*_ to adjust for the system size and to facilitate comparisons with other Kuramoto-model variants. Accordingly, the initial *J*_*ee*_ distribution follows the Gaussian distribution 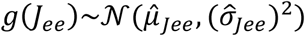, where 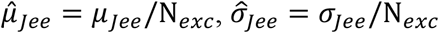. The other interaction strengths, *K*_*ii*_, *K*_*ei*_, and *K*_*ie*_, were uniform and consistently applicable across all units. Equation M5 becomes

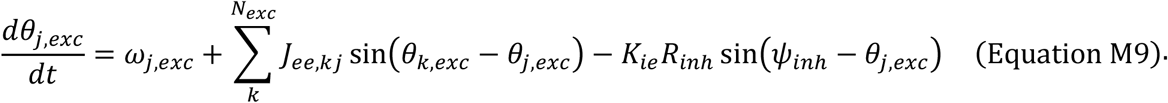

*J*_*ee,kj*_ is the coupling weight between the excUnits *k* (pre) and *j* (post).

Given the rotational symmetry, when the mean natural frequencies of excUnits and inhUnits are equal, 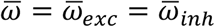, they can be set to 0 rad/s, which is equivalent to rotating the coordinate frame by the mean natural frequency 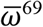. For all simulations in this study: *N*_*exc*_ = 80, *N*_*inh*_ = 20; total simulation duration *T* = 50 s; time step *dt* = 0.01 s. The *g*(*ω*) followed the Gaussian distribution *g*(*ω*)∼𝒩(0, 0.01) for both excUnits and inhUnits. In Figs. 2–5, *K*_*ii*_ and *K*_*ie*_ are unified with the same parameter *K*_*i*_ to simplify the simulation.

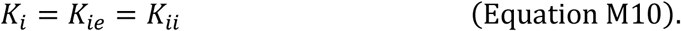

At each time step Δ*t* (= *dt* = 0.01 s in this study), the *J*_*ee*_ values were first modulated by Hebbian potentiation and then rescaled using homeostatic regularization rules. The Hebbian potentiation plasticity is implemented as an exponential function:

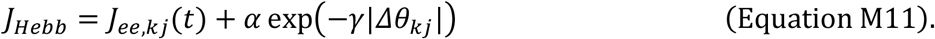

*α* is the learning rate for Hebbian plasticity. Note that *α* should scale proportionally with an inverse of *N*_*exc*_ as α ∝ 1/N_*exc*_ to maintain consistency across different system sizes. *γ* determines the sharpness of the exponential decay from the peak. Δ*θ*_*kj*_ is the phase difference between excUnit *k* and *j*

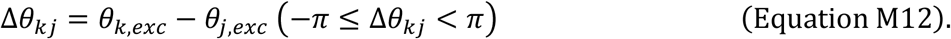

The implementation of Hebbian plasticity with a cosine function (Figure S1A) is defined as

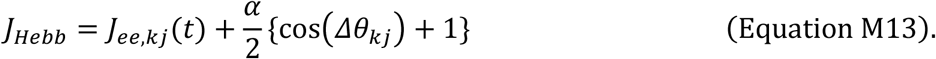

In any Hebbian learning rule in this paper, weight changes of *J*_*ee,kj*_ and *J*_*ee,kj*_ are the same due to the symmetry with respect to the sign of Δ*θ*_*kj*_.

After the Hebbian modulations, *J*_*Hebb*_ was rescaled with the homeostatic regularization plasticity rule to prevent overpotentiation. Plasticity was implemented as follows:

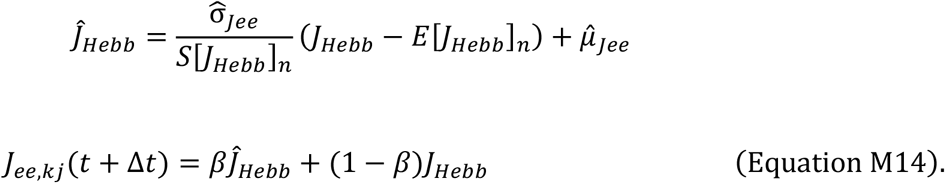

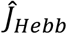 represents the normalized value (homeostatic set-point) of *J*_*Hebb*_ under the initial setting of the mean 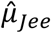 and standard deviation 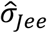. *β* is the learning rate for homeostatic plasticity. This procedure scales *J*_*ee,kj*_ toward the initial targets, 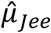 and 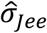.

The change in coupling weight at each time step *ΔJ*_*ee*_ (Figure 3D and E) was defined as

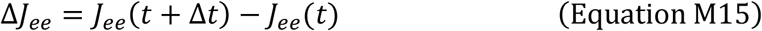

For plotting, pairs of *ΔJ*_*ee*_ and *J*_*ee*_(*t*) values were pooled across all excUnit pairs and time points, and the *J*_*ee*_(t) values were binned (bin size = 0.001).

### Cyclic inhibitory modulation

In Figure 5, *K*_i_ is modulated as a sine function

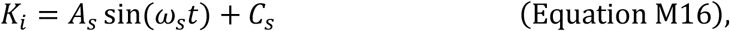

where *A*_*s*_ (= 0.32), *ω*_*s*_ (= 0.004π), and *C*_*s*_ (= 1.28) are the amplitude, frequency, and baseline offset for the sine modulation, respectively. The other parameters for interaction strengths are held constant to simplify the simulation: *μ*_*Jee*_ = 1, *σ*_*Jee*_ = 0.1, *K*_*ei*_ = 3, and *T* = 3,000 s.

### Statistics

E[·]_t_ (also E{·}_t_) and S[·]_t_ are the mean and the standard deviation over time, respectively, where *t* indicates the number of time points. E[·]_n_ and S[·]_n_ are the mean and the standard deviation among unit pairs, respectively, where *n* indicates the number of unit pairs. E[·]_t,n_ and S[·]_t,n_ represent mean and standard deviation, respectively, pooling all values across time and unit pairs. Here, · represents a certain variable. The range of time for the time series analyses (E[·]_t_, S[·]_t_, E[·]_t,n_, and S[·]_t,n_) is [*T*/2, *T*] to exclude the initial fluctuations. For Figure 5B lower panel, value of E{S[*J*_*ee*_]_t_}_n_ was estimated in the time window [-15s, +15 s] for each time point.

Descriptive statistics (mean, standard deviation, kurtosis, and skewness) and statistical tests were performed using the Python NumPy and SciPy libraries. The Dip test^34^ was performed using the Python Diptest Library (https://github.com/RUrlus/diptest). The dip statistic quantifies the maximum distance between the observed distribution and the closest unimodal distribution. If the data distribution is close to unimodal distribution, the dip value is low. If the distribution is bimodal, the dip value is high. Hierarchical clustering was performed using the linkage function of the Python SciPy library.

## Acknowledgments

We would like to thank Susumu Setogawa for his insightful comments on the manuscript. This study was supported by Japan Society for the Promotion of Science KAKENHI [grant 24K18243 to S.K.; grants 23K27479 and 25H02508 to K.M.], Japan Agency for Medical Research and Development (grant JP24wm0625216 to K.M.), and the Takeda Science Foundation [to K.M.]. The funding sources had no involvement in the study design, data collection, interpretation, or the decision to submit this paper for publication.

## Author contributions

Conceptualization: SK; Methodology: SK; Investigation: SK; Visualization: SK; Supervision: KM; Writing—original draft: SK; Writing—review & editing: SK, KM; Funding acquisition: SK, KM

## Data and materials availability

The scripts for this study can be found at Google Colaboratory: https://colab.research.google.com/drive/1v8JF97G9YdwUOh_Wuh9i_TvuGa42fLAu?usp=sharing

