## Supplemental information for "Excitation–inhibition balance controls coupling stability and network reorganization in a plastic Kuramoto model"

**Supplemental Text**

Derivation of the quasi-steady coupling weight distribution

The combined changes from Hebbian plasticity with an exponential function and homeostatic plasticity (Equations M11 and M14) are summarized as follows:

$J_{ee,kj}(t+\Delta t) = J_{ee,kj}(t) + \alpha \exp(-\gamma|\Delta\theta_{kj}|) - \beta\{J_{ee,kj}(t) + \alpha \exp(-\gamma|\Delta\theta_{kj}|) - \hat{J}_{Hebb}\}$ (Equation S1).

Assuming a quasi-steady $J_{ee}$ distribution, $J_{ee,kj}(t+\Delta t) \approx J_{ee,kj}(t)$, if $\alpha$ is small enough, the additive effect of Hebbian potentiation across the entire $J_{ee}$ distribution becomes negligible, resulting in minimal changes to its mean and standard deviation. In this case, the homeostatic set point attained after Hebbian potentiation $\hat{J}_{Hebb}$ can be approximated as that before Hebbian potentiation $\hat{J}_{ee,kj}$, $\hat{J}_{Hebb} \approx \hat{J}_{ee,kj}$. Under these assumptions, the equation simplifies to:

$$\alpha(1-\beta)\exp(-\gamma|\Delta\theta_{kj}|) - \beta(J_{ee,kj} - \hat{J}_{ee,kj}) = 0 \qquad \text{(Equation S2).}$$

Thus,

$$J_{ee,kj} = \hat{J}_{ee,kj} + c\exp(-\gamma|\Delta\theta_{kj}|) \qquad \text{(Equation S3).}$$

$$c = \frac{\alpha(1-\beta)}{\beta}$$

This corresponds to Equation 1 in the main text. The first term, $\hat{J}_{ee,kj}$, represents the homeostatic set point characterized by a $J_{ee}$ distribution with mean $\hat{\mu}_{Jee}$ and variance $\hat{\sigma}_{Jee}$. The second term is the Hebbian potentiation component with a coefficient of $c = \alpha(1-\beta)/\beta$. Assuming a similar process to Equation S3 governs the updates until a quasi-steady state is reached, the $J_{ee}$ distribution sequentially reflects the influence of the $\Delta\theta$-dependent Hebbian term. This indicates that the $\hat{J}_{ee,kj}$ value can also be $\Delta\theta_{kj}$-dependent (Assumption 1). The results of the numerical simulation, which showed that the shapes of $J_{ee}$ and phase difference distributions were correlated (Figure 1E, 1F, and Extended Data Figure 1), supported this assumption.

We defined the probability density function (PDF) of the phases $f_\theta(\theta)$ assuming a sufficiently large number of units:

$$\int_0^{2\pi} f_\theta(\theta)d\theta = 1 \qquad \text{(Equation S4)}.$$

The distribution of the phase difference, $f_{\Delta\theta}(\Delta\theta)$, between two independent phases is given by the circular autocorrelation of the phase distribution:

$$f_{\Delta\theta}(\Delta\theta) = \int_0^{2\pi} f_\theta(\theta) f_\theta(\theta + \Delta\theta)d\theta \ \ (-\pi \leq \Delta\theta < \pi) \qquad \text{(Equation S5)}.$$

In a fully desynchronized state, the phase distribution is uniform, $f(\theta) = 1/2\pi$. In this case, Equation S5 becomes:

$$f_{\Delta\theta}(\Delta\theta) = \int_0^{2\pi} \frac{1}{2\pi} \cdot \frac{1}{2\pi} d\theta = \frac{1}{2\pi} \qquad \text{(Equation S6)}.$$

This shows that the distribution of phase differences is also uniform. Therefore, the absolute phase difference is uniformly distributed over [0,π], $x = |\Delta\theta| \sim \mathrm{Unif}(0, \pi)$, with a PDF:

$$f_{|\Delta\theta|}(x) = \frac{1}{\pi} \qquad \text{(Equation S7)}.$$

Here, we evaluate the mean and standard deviation of the $J_{ee}$ in the quasi-steady state to calculate the homeostatic set point $\hat{J}_{ee,kj}$. We analyze the distribution of the Hebbian component from Equation S3

$$y = c \exp(-\gamma x) \qquad \text{(Equation S8)}.$$

This is a transformation of the variable x ($|\Delta\theta|$). The inverse transformation of Equation S8 is:

$$x = -\frac{1}{\gamma}\log\frac{y}{c} \qquad \text{(Equation S9).}$$

The absolute value of the Jacobian of this transformation is:

$$\left|\frac{dx}{dy}\right| = \frac{1}{\gamma y} \qquad \text{(Equation S10).}$$

We calculate the PDF of the transformed variable $y$, $f_{Jee}(y)$. In the desynchronized state, phase difference distribution follows Equation S7. The relationship between the PDFs is given by

$$f_{|\Delta\theta|}(x)|dx| = f_{Jee}(y)|dy| \qquad \text{(Equation S11).}$$

Substituting Equation S7 and S10 into Equation S11 and using Equation S8 to calculate the range, we obtain:

$$f_{Jee}(y) = f_{|\Delta\theta|}(x)\left|\frac{dx}{dy}\right| = \frac{1}{\gamma\pi y}\ (y_{min} = ce^{-\gamma\pi} \leq y \leq y_{max} = c) \qquad \text{(Equation S12).}$$

Through this variable transformation, the Hebbian component converts the uniform phase difference distribution into a hyperbolic weight distribution, whose probability density is proportional to $1/y$. The mean $E[y]_n$ and variance $Var[y]_n$ of $f_{Jee}(y)$ are:

$$E[y]_n = \int_{y_{min}}^{y_{max}} y f_{Jee}(y)dy = \int_{ce^{-\gamma\pi}}^{c} y\frac{1}{\gamma\pi y}dy = \frac{c(1-e^{-\gamma\pi})}{\gamma\pi} \qquad \text{(Equation S13)}$$

$$\begin{aligned} Var[y]_n &= E[y^2]_n - (E[y]_n)^2 = \int_{ce^{-\gamma\pi}}^{c} y^2\frac{1}{\gamma\pi y}dy - (E[y]_n)^2 \\ &= c^2\left[\frac{1-e^{-2\gamma\pi}}{2\gamma\pi} - \left\{\frac{1-e^{-\gamma\pi}}{\gamma\pi}\right\}^2\right] \end{aligned} \qquad \text{(Equation S14).}$$

Thus, given the standard deviation $S[y]_n = Var[y]_n^{\frac{1}{2}}$, the rescaled $y$ with the homeostatic regulation plasticity, $\hat{y}$, is obtained in a manner similar to that shown in Equation M14 (replacing $J_{Hebb}$ with $y$).

$$\hat{y} = \hat{\sigma}_{Jee}\left(\frac{y - E[y]_n}{S[y]_n}\right) + \hat{\mu}_{Jee}$$

$$\frac{y - E[y]_n}{S[y]_n} = \left(e^{-\gamma|\Delta\theta|} - \frac{1 - e^{-\gamma\pi}}{\gamma\pi}\right) \Big/ \left[\frac{1 - e^{-2\gamma\pi}}{2\gamma\pi} - \left\{\frac{1 - e^{-\gamma\pi}}{\gamma\pi}\right\}^2\right]^{\frac{1}{2}} \qquad \text{(Equation S15).}$$

Thus, $\hat{y}$ is independent of $c$. From Assumption 1, we can consider that the full $J_{ee}$ distribution is dominated by this Hebbian component, allowing us to set $y = J_{ee,kj}$. Therefore, $\hat{y}$ becomes equivalent to the homeostatic set point $\hat{J}_{ee,kj}$. For clarity, $E[y]_n$ and $S[y]_n$ were replaced with the mean and standard deviation of the entire $J_{ee}$ distribution, $E[J_{ee}]_n$ and $S[J_{ee}]_n$. Note that Equation S15 can express the $\hat{J}_{ee,kj}$ as a function of $\Delta\theta_{kj}$. Thus, the value of $\hat{J}_{ee,kj}$ is now calculable.

Next, we consider the change in the phase difference for a pair of excUnits. Given Equation M9 and S3, we calculate $d\Delta\theta_{kj}/dt = d\theta_k/dt - d\theta_j/dt$.

$$\frac{d\Delta\theta_{kj}}{dt} = \Delta\omega - 2\left\{\hat{J}_{ee,kj} + c\exp\left(-\gamma\left|\Delta\theta_{kj}\right|\right)\right\}\sin\left(\Delta\theta_{kj}\right) + F_{exc}^{(-kj)} - F_{inh} \qquad \text{(Equation S16),}$$

where natural frequency difference $\Delta\omega = \omega_k - \omega_j$, repulsive force from the inhUnits $F_{inh} = K_{ie}R_{inh}\{\sin(\psi_{inh} - \theta_k) - \sin(\psi_{inh} - \theta_j)\}$, and attractive force from the excUnits except unit $k$ and $j$, $F_{exc}^{(-kj)}$. Under a mean-field approximation for a sufficiently large number of units, this excitatory force term can be written as:

$$F_{exc}^{(-kj)} = \sum_{l \neq k,j}\left\{J_{ee,kl}\sin(\theta_l - \theta_k) - J_{ee,jl}\sin\left(\theta_l - \theta_j\right)\right\}$$
$$\approx J'^{(-kj)}_{ee} R_{exc}^{(-kj)}\left\{\sin\left(\psi_{exc}^{(-kj)} - \theta_k\right) - \sin\left(\psi_{exc}^{(-kj)} - \theta_j\right)\right\} \qquad \text{(Equation S17),}$$

where $J'^{(-kj)}_{ee}$ represents the effective attractive interaction strength on the pair from other excUnits. $\psi_{exc}^{(-kj)}$ represents the mean phase among excUnits except unit $k$ and $j$. This system also reaches a steady state where $d\Delta\theta_{kj}/dt = 0$. Under this condition,

$$-\Delta\omega + F_{inh} - F_{exc}^{(-kj)} = -2\left\{\hat{J}_{ee,kj} + c\exp\left(-\gamma\left|\Delta\theta_{kj}\right|\right)\right\}\sin\left(\Delta\theta_{kj}\right) \quad \text{(Equation S18)}.$$

This corresponds to Equation 2 in the main text. We can calculate $\hat{J}_{ee,kj}$ with Equation S15. The right-hand side of Equation S18 represents the interaction term between the excUnit $k$ and $j$, while the left-hand side of this equation comprises all other influences, namely the natural frequency difference between the pair and the net mean-field force from the other units. A stable fixed point for $\Delta\theta_{kj}$ exists, where the functions for the left- and right-hand sides intersect (Extended Data Figure 3). In the desynchronized state, the order parameters $R_{exc}$ and $R_{inh}$ are close to zero, limiting the influence of $F_{inh}$ and $F_{exc}^{(-kj)}$. However, in the bistable state, these order parameters fluctuate periodically. Furthermore, the composite sine function $\sin(\psi - \theta_k) - \sin(\psi - \theta_j)$ in $F_{inh}$ and $F_{exc}^{(-kj)}$ is enhanced when the mean phase $\psi$ lies on the shorter arc between $\theta_k$ and $\theta_j$, but is weakened, and can even reverse sign, when it lies on the longer arc. These cause $F_{inh}$ and $F_{exc}^{(-kj)}$ to become influential and to fluctuate in the bistable state; consequently, the $J_{ee}$ distribution is no longer in a quasi-steady state.

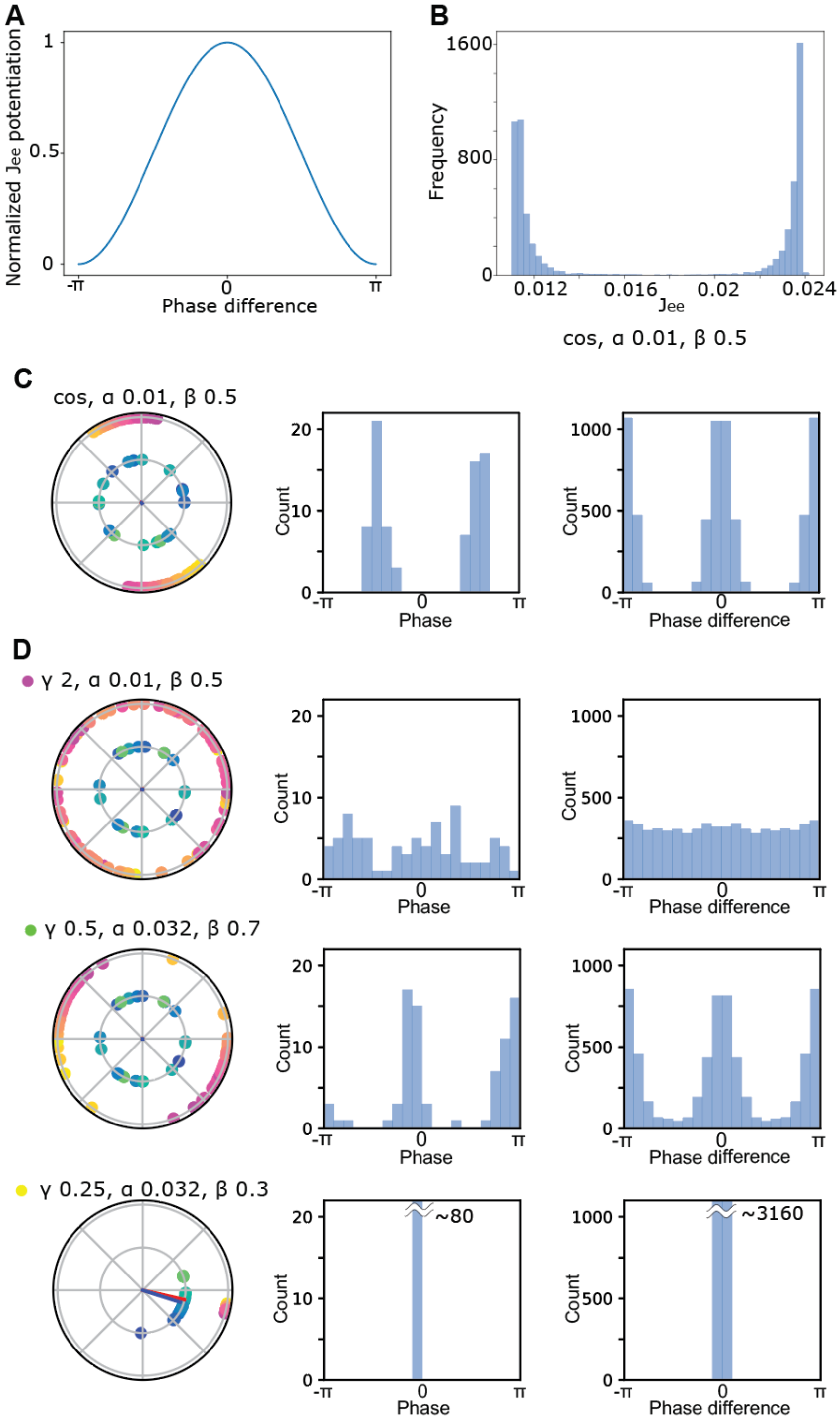


**Figure S1.**

**Cosine Hebbian plasticity rule, phase and phase difference distribution.** (A) Cosine rule for the Hebbian potentiation plasticity. The $J_{ee}$ weight of each pair of units is potentiated based on the phase difference between the units. (B) $J_{ee}$ distribution using the cosine Hebbian plasticity rule at the last frame ($t$ = 50 s). (C) Phase and phase difference distributions at the last frame ($t$ = 50 s) for the cosine plasticity setting. Left panels: Polar plots showing the phases of excUnits (warm colors) and inhUnits (cool colors). Center panels: Histograms of the excUnit phases. Right panels: Histograms of the phase differences between pairs of excUnits. (D) Phase and phase difference distributions at the last frame ($t$ = 50 s) for the exponential plasticity settings. The panel represents the same as (C). The plasticity setting values and circle colors are the same as those in Figure 1E and F. $\mu_{Jee} = 1$, $\sigma_{Jee} = 0.1$, $K_{ei} = 3$, $K_{ie} = 3$, $K_{ii} = 3$.

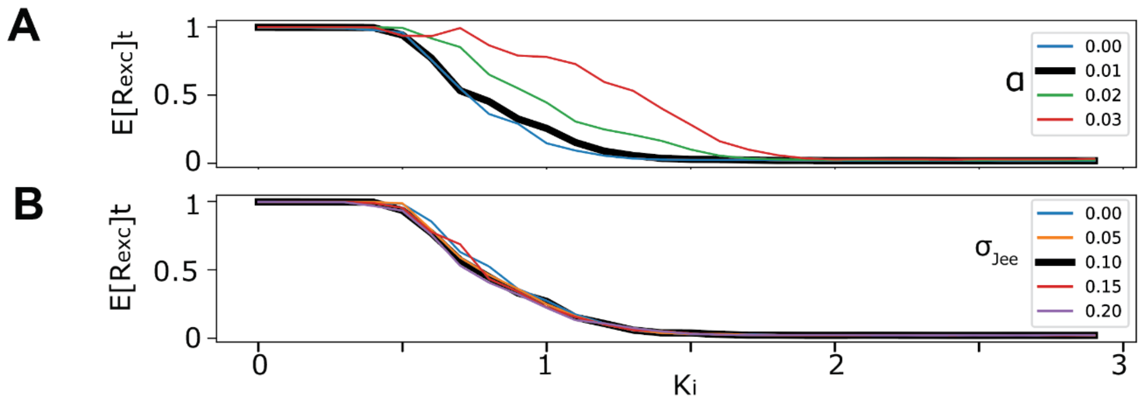


**Figure S2.**

**Effect of exponential Hebbian learning rate and $J_{ee}$ variance on the synchronization.** (A) $E[R_{exc}]_t$ as a function of $K_i$ at different learning rates under Hebbian plasticity ($\alpha$) settings. (B) $E[R_{exc}]_t$ as a function of $K_i$ for different $\sigma_{Jee}$ settings. The thick black lines in (A) and (B) are the same as those in Figure 2C. $\mu_{Jee} = 1$, $K_{ei} = 3$, $\beta = 0.5$, $\gamma = 2$. $\sigma_{Jee} = 0.1$ for (A). $\alpha = 0.01$ for (B).

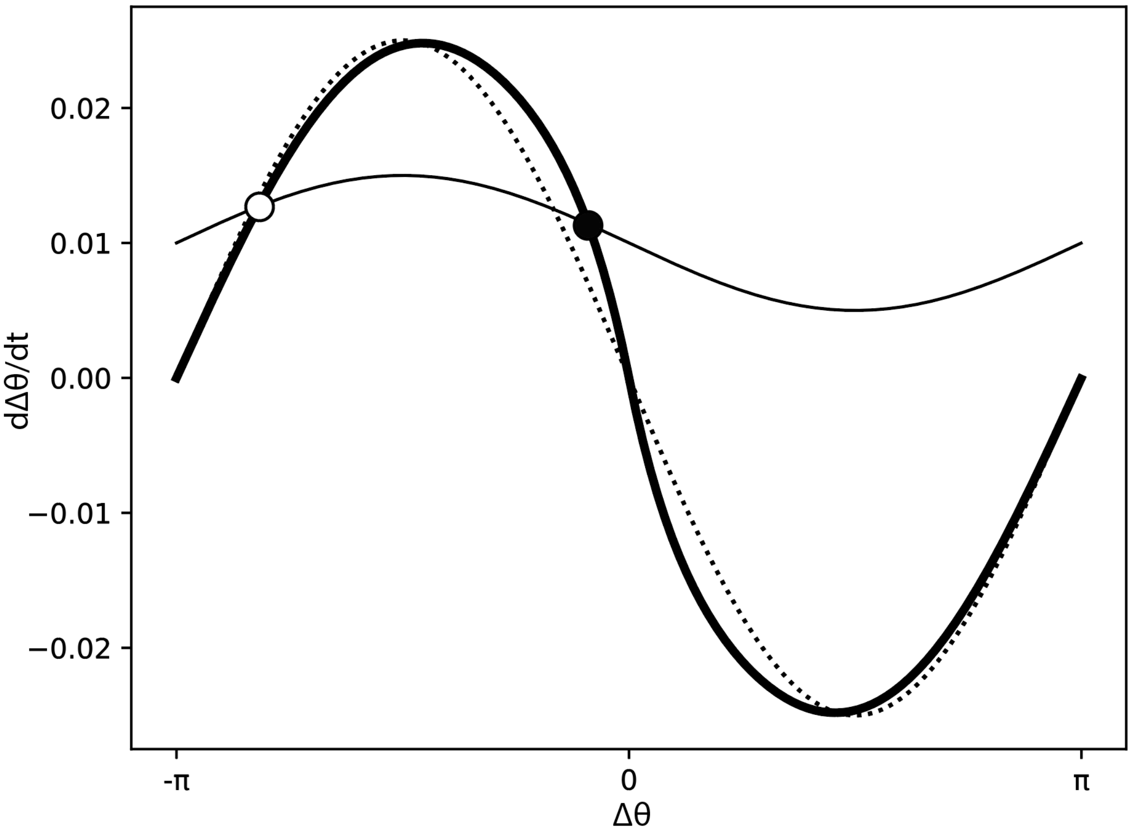


**Figure S3.**

**Interaction function and equilibrium**. The thick solid line indicates the interaction function between excUnits, $-2\left\{\hat{J}_{ee} + c\exp(-\gamma|\Delta\theta|)\right\}\sin(\Delta\theta)$. The thin solid line is the total interaction strength from the other factors, $-\Delta\omega + F_{inh} - F_{exc}^{(-kj)}$. The dotted line indicates the pure sine curve $\hat{\mu}_{Jee}\sin(\Delta\theta)$ for comparison. The black and white points indicate stable and unstable equilibria, respectively. The parameter setting is $\alpha$=0.01, $\beta$=0.5, $\gamma$=2, $\hat{\mu}_{Jee}$ =0.0125 ($\mu_{Jee}/\mathrm{N}_{exc}$= 1/80), $\hat{\sigma}_{Jee}$ = 0.00125 ($\sigma_{Jee}/\mathrm{N}_{exc}$= 0.1/80) (Figure 1F magenta). In this figure, as an example, we set $\Delta\omega$ = -0.01, $F_{inh} - F_{exc}^{(-kj)} = -0.2\sin\Delta\theta$.

**Video S1.**

Synchronized state: $\mu_{Jee} = 1$, $\sigma_{Jee} = 0.1$, $K_{ei} = 3$, $K_i = 0.1$, $\gamma = 2$, $\alpha = 0.01$, $\beta = 0.5$; playback speed: ×1

**Video S2.**

Desynchronized state: $\mu_{Jee} = 1$, $\sigma_{Jee} = 0.1$, $K_{ei} = 3$, $K_i = 1.5$, $\gamma = 2$, $\alpha = 0.01$, $\beta = 0.5$; playback speed: ×1.

**Video S3.**

Bistable state 1: $\mu_{Jee} = 1$, $\sigma_{Jee} = 0.1$, $K_{ei} = 3$, $K_i = 0.8$, $\gamma = 2$, $\alpha = 0.01$, $\beta = 0.5$; playback speed: ×1.

**Video S4.**

Bistable state 2: $\mu_{Jee} = 1$, $\sigma_{Jee} = 0.1$, $K_{ei} = 1$, $K_i = 1$, $\gamma = 2$, $\alpha = 0.01$, $\beta = 0.5$; playback speed: ×1.

**Video S5.**

Cyclic inhibitory modulation of Ki; playback speed ×20.
